# Adaptive trends of sequence compositional complexity over pandemic time in the SARS-CoV-2 coronavirus

**DOI:** 10.1101/2021.11.06.467547

**Authors:** José L. Oliver, Pedro Bernaola-Galván, Francisco Perfectti, Cristina Gómez-Martín, Silvia Castiglione, Pasquale Raia, Miguel Verdú, Andrés Moya

## Abstract

During the spread of the COVID-19 pandemic, the SARS-CoV-2 coronavirus underwent mutation and recombination events that altered its genome compositional structure, thus providing an unprecedented opportunity to search for adaptive evolutionary trends in real-time. The mutation rate in coronavirus is known to be lower than expected for neutral evolution, thus suggesting a role for natural selection. We summarize the compositional heterogeneity of each viral genome by computing its Sequence Compositional Complexity (SCC). To study the full range of SCC diversity, random samples of high-quality coronavirus genomes covering pandemic time span were analyzed. We then search for evolutionary trends that could inform on the adaptive process of the virus to its human host by computing the phylogenetic ridge regression of SCC against time (i.e., the collection date of each viral isolate). In early samples, we find no statistical support for any trend in SCC, although the viral genome appears to evolve faster than Brownian Motion (BM) expectation. However, in samples taken after the emergence of high fitness variants, and despite the brief time span elapsed, a driven decreasing trend for SCC, and an increasing one for its absolute evolutionary rate, are detected, pointing to a role for selection in the evolution of SCC in coronavirus genomes. We conclude that the higher fitness of variant genomes leads to adaptive trends of SCC over pandemic time in the coronavirus.

## Introduction

Given the difficulties to observe evolution over the human life scale, test tube experiments revealed as a particularly powerful tool for examining evolutionary dynamics. Richard Lenski’s long-term evolution experiment (LTEE) with a laboratory population of *Escherichia coli* sampled through 60,000 generations shows the relationships between rates of genomic evolution and organismal adaptation (Barrick *et al*., 2009; Good *et al*., 2017). Experimental evolution of a major evolutionary innovation (the origin of multicellularity) has been carried out on both experimentally tractable model organisms (Ratcliff *et al*., 2012), as well as in a unicellular relative of animals (Burnetti and Ratcliff, 2022). In the same way, computer simulations of digital organisms revealed important aspects of evolutionary dynamics (Adami, Ofria and Collier, 2000). Now, the outbreak of the COVID-19 pandemic provides an unprecedented opportunity to observe evolutionary trends in real time, which could provide helpful information on the adaptive process of the viral genome to its human host.

Pioneering works showed that RNA viruses are excellent material for studies of evolutionary genomics (Domingo, Webster and Holland, 1999; Worobey and Holmes, 1999; Moya, Holmes and González-Candelas, 2004). Despite the difficulties of inferring reliable phylogenies of SARS-CoV-2 (Morel *et al*., 2020; Pipes *et al*., 2021), as well as the controversy surrounding the first days and location of the pandemic (Koopmans *et al*., 2021; Worobey, 2021), the most parsimonious explanation for the origin of SARS-CoV-2 seems to lie in a zoonotic event (Zhang and Holmes, 2020; Holmes *et al*., 2021; Balloux *et al*., 2022). Despite its proofreading mechanism and the brief time-lapse since its appearance, SARS-CoV-2 has accumulated an important amount of genomic and genetic variability (Elbe and Buckland-Merrett, 2017; Hatcher *et al*., 2017; Hadfield *et al*., 2018; Dorp *et al*., 2020; Islam *et al*., 2020; Hamed *et al*., 2021; McBroome *et al*., 2021), dramatically impacting viral nucleotide composition and genome organization. Synonymous and non-synonymous mutations (Banerjee *et al*., 2020; Cai, Cai and Li, 2020; González-Candelas *et al*., 2021), as well as mismatches and deletions in translated and untranslated regions (Islam *et al*., 2020; Young *et al*., 2020) have been tracked in the SARs-CoV-2 genome. This may be related to both its recombinational origin (Naqvi *et al*., 2020) as well as mutation and additional recombination events accumulated later (Cyranoski, 2020; Jackson *et al*., 2021).

Recent phylogenetic estimates of the substitution rate of SARS-CoV-2 suggest that its genome accumulates around two mutations per month. However, Variants of Concern (VoCs) can have many more defining mutations (Neher, 2022), and it is hypothesized that they emerged over the course of a few months, implying that they must have evolved faster for a period of time (Tay *et al*., 2022). Noteworthy, RNA viruses can also accumulate high genetic variation during individual outbreaks (Pybus, Tatem and Lemey, 2015), showing mutation and evolutionary rates up to a million times higher than those of their hosts (Islam *et al*., 2020).

Particularly interesting are those changes increasing viral fitness (Dorp *et al*., 2020; Garvin *et al*., 2020; Zhou *et al*., 2020; Holmes *et al*., 2021), such as mutations giving rise to epitope loss and antibody escape mechanisms. These have mainly been found in SARS-CoV-2 variants isolated from Europe and the Americas, and have critical implications for the virus fitness (transmission, pathogenesis, and immune interventions (Gupta and Mandal, 2020; Loucera *et al*., 2022)). Some studies (Neher, 2022), though, have shown that SARS-CoV-2 is acquiring mutations more slowly than expected for neutral evolution, suggesting the action of natural selection. Parallel mutations in multiple independent lineages and variants have been observed (Dorp *et al*., 2020), which may indicate convergent evolution (Cooper, 2021), and are of particular interest in the context of adaptation of SARS-CoV-2 to the human host. Survival analysis of mutations in the SARS-CoV-2 genome revealed 27 of them were significantly associated with higher mortality of patients (Loucera *et al*., 2022). Other authors have reported some sites under positive selection pressure in the nucleocapsid and spike genes (Benvenuto *et al*., 2020). This impressive research effort has allowed tracking all these changes in realtime. The CoVizu^e^ project (https://filogeneti.ca/covizu/) provides a near real-time visualization of SARS-CoV-2 global diversity, the COVID-19 CG website (Chen *et al*., 2021) tracks SARS-CoV-2 mutation and lineage by locations and dates of interest, while the CoV-Spectrum website (Chen *et al*., 2022) supports the identification of new SARS-CoV-2 variants of concern and the real-time tracking of known variants. Another recent tool (Sanderson, 2022) allows a visualization of mutation-annotated trees of millions SARS-CoV-2 sequences (https://cov2tree.org/).

Nucleotide compositional biases throughout the genome have been identified at all levels of the phylogenetic hierarchy (Filipski, Thiery and Bernardi, 1973; Macaya, Thiery and Bernardi, 1976; Bernardi *et al*., 1985; Bernardi, 1990, 2004; Sueoka, 1992; Belle, Smith and Eyre-Walker, 2002; Belle *et al*., 2004; Bernaola-Galván *et al*., 2004; Muto and Osawa, 2006; Moya *et al*., 2020), including RNA virus (Gaunt and Digard, 2022), being caused either by active selection or passive mutation pressure (Mooers and Holmes, 2000). The array of compositional domains in a genome can be potentially altered by most sequence changes (i.e., synonymous and non-synonymous nucleotide substitutions, insertions, deletions, recombination events, chromosome rearrangements, or genome reorganizations). Compositional domain structure can be altered either by changing nucleotide frequencies in a given region or by changing the nucleotides at the borders separating two domains, thus enlarging/shortening a given domain, or changing the number of domains (Bernaola-Galván, Román-Roldán and Oliver, 1996; Oliver *et al*., 1999; Wen and Zhang, 2003; Keith, 2008). Ideally, a metric of nucleotide compositional heterogeneity should be able to summarize all the mutational and recombinational events accumulated by a genome sequence over time (Román-Roldán, Bernaola-Galván and Oliver, 1998; Bernaola-Galván *et al*., 2004; Fearnhead and Vasilieou, 2009).

In many organisms, the patchy sequence structure formed by the array of compositional domains with different nucleotide composition (i.e., GC content) has been related to important biological features, as gene and repeat densities, timing of gene expression, recombination frequency, etc. (Bernardi *et al*., 1985; Oliver *et al*., 2004; Bernaola-Galván, Carpena and Oliver, 2008; Bernardi, 2015). Therefore, changes in sequence compositional heterogeneity may be relevant on evolutionary and epidemiological grounds. Specifically, the existence of evolutionary trends in the compositional complexity of the coronavirus could reveal whether natural selection is providing adaptation of the virus to the human host.

To search for such trends, we computed the Sequence Compositional Complexity, or *SCC* (Román-Roldán, Bernaola-Galván and Oliver, 1998), an entropic measure of nucleotide compositional heterogeneity, representing the number of domains and nucleotide differences among them, identified in a viral genome sequence through a proper segmentation algorithm (Bernaola-Galván, Román-Roldán and Oliver, 1996). The samples of high-quality coronavirus genomes analyzed here show sufficient variation in SCC to derive its genealogical or evolutionary relationships. By using phylogenetic ridge regression, a method able to reveal both macro- (Melchionna *et al*., 2019; Serio *et al*., 2019) and micro-evolutionary (Moya *et al*., 2020) trends, we present here evidence for long-term adaptive tendencies of decreasing sequence compositional heterogeneity, and an increasing one for its evolutionary rate, in SARS-CoV-2. Both trends are shared by its most important VoCs (Alpha and Delta), being greatly accelerated by the recent rise to dominance of Omicron (Du, Gao and Wang, 2022).

## Results

### Sequence compositional complexity in the coronavirus

The reference SARS-CoV-2 coronavirus genome sequence (hCoV-19/Wuhan/WIV04/2019|EPI_ISL_402124|2019-12-30) was divided into eight compositional domains by the compositional segmentation algorithm (Bernaola-Galván, Román-Roldán and Oliver, 1996; Oliver *et al*., 1999, 2004; Bernaola-Galván, Carpena and Oliver, 2008), resulting in a *SCC* value of 5.7 × 10E-3 bits by sequence position (Figure 1).

**Figure 1:**
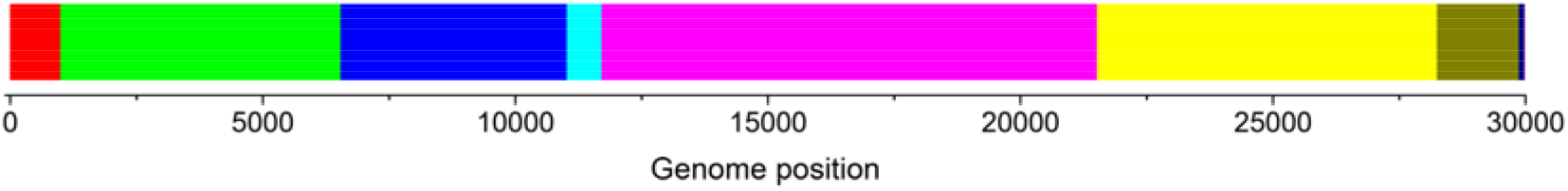
Compositional segmentation of the GISAID reference genome (available at https://www.gisaid.org/resources/hcov-19-reference-sequence/). Using an iterative segmentation algorithm (Bernaola-Galván, Román-Roldán and Oliver, 1996; Oliver *et al*., 2004), and a threshold P value ≤ 0.05, the RNA sequence was divided into eight compositionally homogeneous segments (i.e., compositional domains). The location of domain borders is shown on the horizontal scale. Colors are used only to illustrate the differential nucleotide composition of each domain.

From then on, descendent coronaviruses presented substantial variation in each domain’s number, length, and nucleotide composition, which is reflected in a variety of *SCC* values. The number of segments ranges between 4 and 10, while the *SCC* do so between 2.71E-03 and 6.8E-03 bits by sequence position. Therefore, coronavirus genomes show sufficient variation in SCC to derive its genealogical or evolutionary relationships. The strain name, the collection date, and the *SCC* values for each analyzed genome are shown in Supplementary Tables S1-S19 available in the open repository Zenodo (https://doi.org/10.5281/zenodo.7555406).

### Temporal evolution of SCC over the coronavirus pandemic

To characterize the temporal evolution of *SCC* over the coronavirus pandemic, we downloaded from GISAID/Audacity (Elbe and Buckland-Merrett, 2017; Shu and McCauley, 2017; Khare *et al*., 2021) a series of random samples of high-quality genome sequences over consecutive time lapses, each starting at the outbreak of the COVID-19 (December 2019) and progressively including younger samples up to October 2022 (Table 1). In each random sample, we filtered and masked the genome sequences using the GenBank reference genome MN908947.3 to eliminate sequence oddities (Hodcroft, Domman, *et al*., 2021). Non-duplicated genome sequences were aligned with *MAFFT* (Katoh and Standley, 2013), then inferring the best ML tree using *IQ-TREE 2* (Minh *et al*., 2020). Using the least square dating (LSD2) method (To *et al*., 2016) a timetree was build, which was finally rooted to the GISAID reference genome (hCoV-19/Wuhan/WIV04/2019|EPI_ISL_402124|2019-12-30). The proportion of variant genomes in each sample was determined with *Nextclade* (Aksamentov *et al*., 2021) (Table 1, columns 5-8).

**Table 1.** Phylogenetic trends in random coronavirus samples downloaded from the GISAID database Audacity (Elbe and Buckland-Merrett, 2017; Shu and McCauley, 2017; Khare *et al*., 2021) covering the pandemic time range from December 2019 to March 2022. For each sample, the analyzed time range was from December 2019 to the date shown in the column ‘Collection date’. Initial sample sizes were 500, 1,000, 2,000, or 3,000 genomes per sample, while the final sample size indicates the remaining genome sequences once duplicated sequences were discarded. Non-duplicated genomes in each sample were then aligned with *MAFFT* (Katoh and Standley, 2013) to the GenBank MN908947.3 reference genome sequence and masked to eliminate sequence oddities (Hodcroft, De Maio, *et al*., 2021). The best ML timetree for each sample was inferred using IQ-TREE 2 (Minh *et al*., 2020), which was rooted to the GISAID reference genome (hCoV-19/Wuhan/WIV04/2019|EPI_ISL_402124|2019-12-30). The percentages of variant genomes were determined by *Nextclade* (Aksamentov *et al*., 2021). The genome heterogeneity of each coronavirus genome was determined by computing its Sequence Compositional Complexity, or SCC (Román-Roldán, Bernaola-Galván and Oliver, 1998). Phylogenetic ridge regressions for SCC and its evolutionary rate were computed by the function *search.trend* from the *RRphylo* R package (Castiglione *et al*., 2018). The estimated genomic value for each tip or node in the phylogenetic tree is regressed against time. The statistical significance of the ridge regression slope was tested against 1,000 slopes obtained after simulating a simple (i.e., no-trend) Brownian evolution of SCC in the phylogenetic tree. See Methods for further details.

|  |  | Sample size |  | % Of main variants |  |  | Total of variants in the sample (%) | SCC regression |  | Rate regression |  |
| --- | --- | --- | --- | --- | --- | --- | --- | --- | --- | --- | --- |
| Sample | Collection date | Initial | Final | Alpha | Delta | Ómicron |  | Slope | P value | Slope | P value |
| s726 | mar-20 | 1,000 | 726 | 0.00 | 0.00 | 0.00 | 0.00 | -97.10 | 0.250 | 88,684.70 | 0.002 |
| s730 |  | 1,000 | 730 | 0.00 | 0.00 | 0.00 | 0.00 | -65.20 | 0.268 | 145,566.20 | 0.001 |
| s781 |  | 2,000 | 781 | 0.00 | 0.00 | 0.00 | 0.00 | -9.67 | 0.426 | 125,140.00 | 0.001 |
| s1170 | jun-20 | 2,000 | 1,170 | 0.00 | 0.00 | 0.00 | 0.00 | 12.83 | 0.444 | 87,637.54 | 0.001 |
| s1277 | sep-20 | 2,000 | 1,277 | 0.00 | 0.00 | 0.00 | 0.00 | -20.85 | 0.305 | 39,183.10 | 0.001 |
| s1573 | dec-20 | 2,000 | 1,573 | 4.32 | 0.00 | 0.00 | 4.83 | -38.53 | 0.066 | 26,502.59 | 0.001 |
| s1871 | mar-21 | 2,000 | 1,871 | 50.03 | 0.00 | 0.00 | 57.03 | -61.94 | 0.001 | 14,254.56 | 0.001 |
| s498 | may-21 | 500 | 498 | 56.43 | 0.00 | 0.00 | 64.65 | -66.39 | 0.011 | 15,035.58 | 0.001 |
| s496 |  | 500 | 496 | 64.52 | 1.41 | 0.00 | 73.79 | -55.74 | 0.026 | 11,090.87 | 0.001 |
| s987 |  | 1,000 | 987 | 57.85 | 0.20 | 0.00 | 67.17 | -60.29 | 0.001 | 16,937.26 | 0.001 |
| s980 |  | 1,000 | 980 | 65.31 | 0.82 | 0.00 | 75.41 | -54.23 | 0.004 | 16,169.35 | 0.001 |
| s1939 |  | 2,000 | 1,939 | 63.02 | 1.24 | 0.00 | 72.93 | -41.74 | 0.010 | 13,044.29 | 0.001 |
| s1974 | jun-21 | 2,000 | 1,974 | 45.90 | 7.14 | 0.00 | 60.59 | -34.83 | 0.016 | 18,624.45 | 0.001 |
| s1985 | sep-21 | 2,000 | 1,985 | 27.10 | 44.08 | 0.00 | 78.34 | -19.30 | 0.131 | 11,688.97 | 0.001 |
| s1994 | dec-21 | 2,000 | 1,994 | 17.95 | 57.82 | 6.22 | 86.70 | -20.93 | 0.060 | 7,495.37 | 0.001 |
| s2347 |  | 3,000 | 2,347 | 18.41 | 46.66 | 0.00 | 73.11 | -33.38 | 0.007 | 7,217.62 | 0.001 |
| s1990 | mar-22 | 2,000 | 1,990 | 14.32 | 51.41 | 18.29 | 87.28 | -21.89 | 0.037 | 4,896.06 | 0.052 |
| s3037 | oct-22 | 3,060 | 3,037 | 8.99 | 30.52 | 50.38 | 92.10 | -42.26 | 0.001 | 6,111.83 | 0.001 |
| TOTAL: |  | 31,060 | 26,355 |  |  |  |  |  |  |  |  |

We sought temporal phylogenetic trends in *SCC* values and evolutionary rates by using the function *search.trend* in the *RRphylo* R package (Castiglione *et al*., 2018), contrasting the realized slope of *SCC* versus time regression to a family of 1,000 slopes generated under the BM model of evolution, which models evolution with no trend in either the *SCC* or its evolutionary rate. We found that SARS-CoV-2 sequence compositional heterogeneity did not follow any trend in *SCC* during the first year of the pandemic time course, as indicated by the non-significant *SCC* against time regressions in any sample ending before December 2020 (Table 1). However, with the emergence of variants in December 2020, the SCC vs time slope started to decrease significantly. But in contrast to the decreasing trend observed for *SCC*, a clear tendency towards faster evolutionary rates occurred throughout the study period, indicating that the virus increased in variability early on, but took on a monotonic trend of fast decreasing *SCC* as VoCs appeared. These results were robust to several sources of uncertainty, including those related to the algorithms used for multiple alignment or to infer phylogenetic trees (see the section ‘Checking results reliability’ in Supplementary Information) and phylogenetic uncertainty itself. A strong acceleration of the overall rate compared to within clade evolution has been previously observed by estimating the rates of evolution for synonymous and non-synonymous changes separately for evolution within clades and for the pandemic overall (Neher, 2022), indicating that the evolutionary process that gave rise to the different variants is qualitatively different from that in typical transmission chains, being likely dominated by adaptive evolution. In summary, our analyses show that statistically significant trends for declining SCC began between the end of December 2020 (s1573) and March 2021 (s1871) corresponding with the emergence of the first VoC (Alpha), a path that continued with the successive emergence of other variants. This may suggest a role for natural selection in the evolution of SCC in the coronavirus. In fact, in a parasite/host system, we should expect episodic natural selection and some fluctuating selection.

### Relative contributions of individual variants to the SARS-CoV-2 evolutionary trends

#### SCC trends of individual variants of concern

We estimated the relative contribution of the three main VoCs (Alpha, Delta, and Omicron) to the trends in SARS-CoV2 evolution by picking samples both before (s726, s730) and after (s1871, s1990, s3037) their appearance. The trends for *SCC* and its evolutionary rate in sample s3037, which includes a sizeable number of Omicron genomes, are shown in Figure 2. For all these samples, we tested trends for variants individually (as well as for the samples’ trees as a whole) while accounting for phylogenetic uncertainty, by randomly altering the phylogenetic topology and branch lengths 100 times per sample (see Methods and Supplementary Information for details). These precautions were taken to ensure accuracy in the conclusions based on the SARs-CoV-2 phylogenies we inferred (Wertheim, Steel and Sanderson, 2022). In agreement with the previous analyses (eighteen consecutive bins, see Table 1), we found strong support for a decrease in *SCC* values through time along phylogenies including variants (s1871, s1990, s3037) and no support for any temporal trend in older samples. Just four out of the 200 random trees produced for samples s726 and s730 produced a trend in *SCC* evolution. The corresponding figure for the two younger samples (s1871, s1990) is 186/200 significant and negative instances of declining *SCC* over time (Table 2). This ~50-fold increase in the likelihood of finding a consistent trend in declining *SCC* over time is shared unambiguously by all tested variants (Alpha, Delta, and Omicron; Table 3). Yet, Omicron shows a significantly stronger decline in *SCC* than the other variants (Table 3), suggesting that the trends starting with the appearance of the main variants became stronger with the emergence of Omicron by the end of 2021.

**Figure 2:**
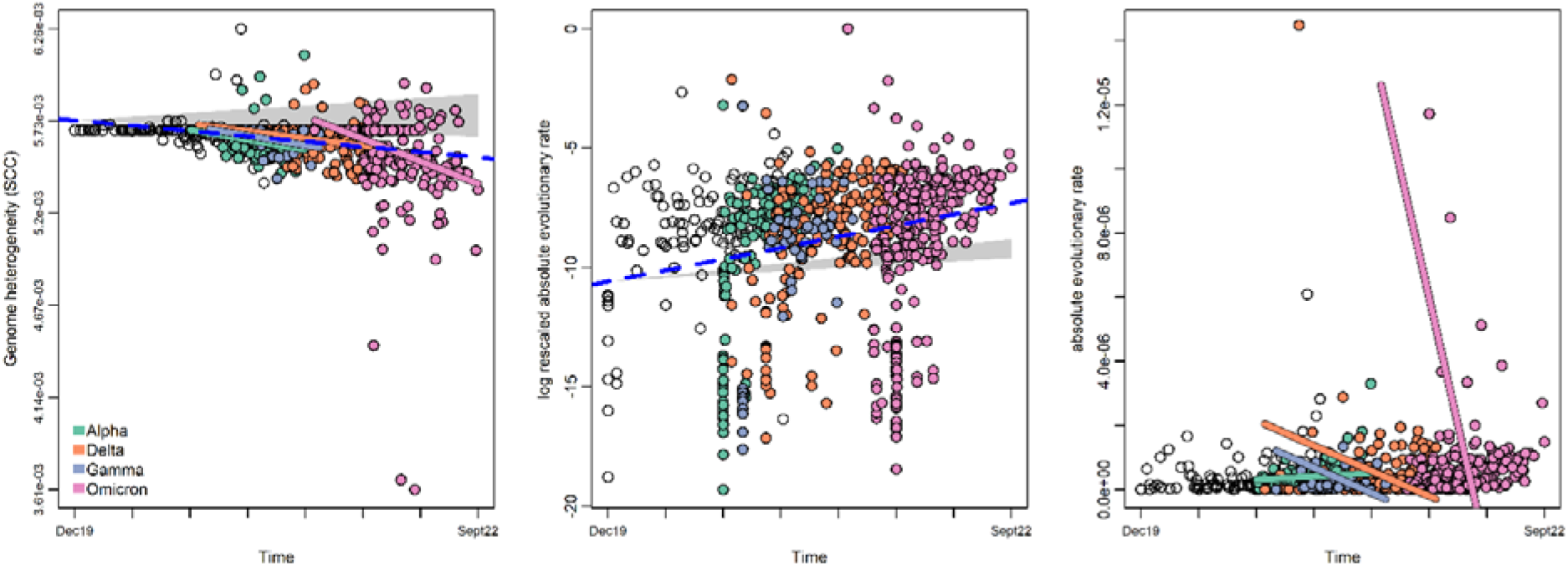
Phylogenetic ridge regressions for *SCC* (left), the rescaled absolute evolutionary rate (center) and the absolute evolutionary rate (right) as detected by the *RRphylo* R package (Castiglione *et al*., 2018) on the s3037 sample. For *SCC* values (left), the estimated value at each node in the tree and the observed tip values are regressed against their age (the phylogenetic time distance, meant mainly as the collection date of each virus isolate and node ages). The dashed blue line represents the regression line obtained for the whole tree, regardless of variants identity. The rescaled evolutionary rate (center) was obtained by rescaling the absolute rate in the 0-1 range and then transforming to logs to compare to the BM expectation (shaded gray area). Eventually, the plot to the right refer to the absolute rate values (unscaled) plotted against time. The statistical significance of the ridge regression slopes was tested against 1,000 slopes obtained after simulating a simple Brownian evolution of the *SCC* in the phylogenetic tree. The 95% confidence intervals around each point produced according to the BM model of evolution are shown as shaded areas. Dots are colored according to the variant they belong to or left blank for strains collected before the appearance of variants.

**Table 2.** Percentages of significant results of SCC and SCC evolutionary rates *versus* time regressions performed on 100 randomly fixed (and subsampled for s1871 and s1990) phylogenetic trees. higher/lower than BM = the percentage of simulation producing slopes significantly higher/lower than the Brownian Motion expectation.

| Sample | SCC values |  | SCC evolutionary rates |  |
| --- | --- | --- | --- | --- |
|  | positive | negative | positive | negative |
| s726 | 0 | 4 | 88 | 0 |
| s730 | 0 | 0 | 100 | 0 |
| s1871 | 0 | 100 | 38 | 0 |
| s1990 | 0 | 86 | 100 | 0 |
| s3037 | 0 | 99 | 100 | 0 |

**Table 3.** Percentages of significant results of SCC and SCC evolutionary rates versus time regressions performed on 100 randomly resolved (s1871 and s1990) phylogenetic trees. % Slope difference indicates the percentage of simulations producing significantly higher/lower slopes than the rest of the tree.

| Sample | Variant |  | SCC values | SCC evolutionary rates |
| --- | --- | --- | --- | --- |
|  |  |  | % Slope difference | % Slope difference |
| s1871 | Alpha | positive | 0 | 0 |
|  |  | negative | 99 | 39 |
| s1990 | Alpha | positive | 0 | 0 |
|  |  | negative | 100 | 0 |
|  | Delta | positive | 0 | 0 |
|  |  | negative | 91 | 27 |
|  | Ómicron | positive | 0 | 0 |
|  |  | negative | 94 | 100 |
| s3037 | Alpha | positive | 0 | 0 |
|  |  | negative | 82 | 0 |
|  | Delta | positive | 0 | 0 |
|  |  | negative | 70 | 0 |
|  | Ómicron | positive | 0 | 0 |
|  |  | negative | 92 | 100 |

We tested the difference in the slopes of *SCC* values versus time regression computed by grouping all the variants under a single group and the same figure for all other strains grouped together. The test was performed using the function *emtrends* available within the R package *emmeans* (Lenth, 2022). We found the slope for the group that includes all variants to be significantly larger than the slope for the other strains (estimate = −0.772 × 10-8, P value = 0.006), still pointing to the decisive effect of VoCs on SCC temporal trend.

#### SCC evolutionary rates of variants

*SCC* evolutionary rate (absolute magnitude of the rate) tends to increase over time (Table 2). The slope of *SCC* rates through time regression for Omicron was always significantly lower than the slope computed for the rest of the tree (Table 3). This was also true for Alpha and Delta, although with much lower support.

## Discussion

Here we show that despite its short length (29,912 bp for the reference genome) and the brief time-lapse analyzed, the coronavirus RNA genome sequences are segmented (Fig. 1 and Supplementary Tables S1-S19) into 4 to 10 compositional domains (~0.27 segments by kbp on average). Although such segment density is lower than in free-living organisms, like cyanobacteria where an average density of 0.47 segments by kbp was observed (Moya *et al*., 2020), the large variability found in different coronavirus strains may suffice for comparative evolutionary analyses of SCC in these genomes, which might shed light on the origin and evolution of the COVID-19 pandemic.

In early samples (i.e., collected before the emergence of variants), we found no statistical support for any trend in *SCC* values over time, although the virus as a whole appears to evolve faster than BM expectation (that is no acceleration or deceleration either). However, save s1985, in the remaining samples taken after the first higher fitness VoC with higher transmissibility (Alpha) appeared in the GISAID database (December 2020), we started to detect statistically significant downward trends in *SCC* (Table 1). Concomitantly to the temporal decay in *SCC*, its absolute evolutionary rate kept increasing with time, meaning that the decline in *SCC* itself accelerated over time. In agreement with this notion, although declining *SCC* is an evolutionary path shared by variants, the nearly threefold increase in rates intensified after the appearance of the most recent VoC (Omicron) in late 2021, which shows a much faster decline in *SCC* than the other variants (Table 3). These results indicate the existence of a driven, most probably adaptive, trend in the variants toward a reduction of SCC.

Adaptive mutations in coronavirus VOCs are well documented in the literature. The emergence of VOCs has been associated to an episodic increase in the substitution rate of around 4-fold the background phylogenetic rate estimate (Tay *et al*., 2022). It is also known that variant genomes have accumulated a higher proportion of adaptive mutations, which allows them to neutralize host resistance or escape host antibodies (Mlcochova *et al*., 2021; Thorne *et al*., 2021; Venkatakrishnan *et al*., 2021), consequently gaining higher transmissibility (a paradigmatic example is the recent outbreak of the Omicron variant). The sudden increases in fitness of variant genomes, may be due to the gathering of co-mutations that become prevalent world-wide compared to single mutations, being largely responsible for their temporal changes in transmissibility and virulence (Ilmjärv *et al*., 2021; Majumdar and Niyogi, 2021). In fact, more contagious and perhaps more virulent VoCs share mutations and deletions that have arisen recurrently in distinct genetic backgrounds (Richard *et al*., 2021). Other authors have reported that some SARS-CoV-2 variants share similar combinations of mutations (Cooper, 2021), also pointing to the existence of convergent evolution in the coronavirus. In this way, the association we observed between SCC and fitness may be explained by the accumulation of different classes of mutations (Benvenuto *et al*., 2020; Korber *et al*., 2020) in emergent variants: main, higher-fitness mutations (i.e., located at the nucleocapsid or the spike genes, which serve to variant definition) and hitchhiking mutations (i.e., located at other genome regions). Homology modeling have also revealed some structural differences between the viruses (Benvenuto *et al*., 2020), which may be also lead to SCC changes. The accumulation of all these point and structural mutational changes may have leads to a lower genome heterogeneity and, consequently, to the SCC decreases we observed.

In conclusion, we show here that increases in fitness of variant genomes, associated with a higher transmissibility, lead to a reduction of their sequence compositional heterogeneity, thus explaining the general decay of *SCC* in line with the pandemic expansion. The accelerated loss of SCC in the coronavirus may be promoted by both the rise of high viral fitness variants and convergent evolution, leading to adaptation to the human host, a well-known process in other viruses (Bahir *et al*., 2009). Further monitoring of the evolutionary trends in current and new co-mutations, variants, and recombinant lineages (Callaway, 2022; Ledford, 2022; Straten *et al*., 2022) by means of the tools used here will enable to elucidate whether and to what extent the evolution of SCC in the virus impacts human health.

## Methods

### Data retrieval, filtering, masking and alignment

The sequences of the high-quality coronavirus genomes retrieved from the GISAID/Audacity database (Elbe and Buckland-Merrett, 2017; Shu and McCauley, 2017; Khare *et al*., 2021) were compiled as EPI_SET_230117nx, being available at https://doi.org/10.55876/gis8.230117nx. *MAFFT* (Katoh and Standley, 2013) was used to align each random sample to the genome sequence of the isolate Wuhan-Hu-1 (GenBank accession MN908947.3), then filtering and masking the alignments to avoid sequence oddities (Hodcroft, Domman, *et al*., 2021). In order to check the reliability of our results (see the section ‘Checking results reliability’ in Supplementary Information), we also analyzed other 3,059 genomes of the SARS-CoV-2 *Nextstrain* global dataset (Hadfield *et al*., 2018) downloaded from https://nextstrain.org/ncov/open/global?f_host=Homo%20sapiens on 2021-10-08. The sequences for this dataset (in Fasta format) and the ML phylodynamic tree obtained by these authors by means of the TreeTime software (Sagulenko, Puller and Neher, 2018) are available at Zenodo (https://doi.org/10.5281/zenodo.7555406).

### Phylogenetic trees

The best ML tree for each random sample in Table 1 was inferred using *IQ-TREE 2* (Minh *et al*., 2020), using the GTR nucleotide substitution model (Tavaré, 1986; Rodríguez *et al*., 1990). We then used_the least square dating (LSD2) method (To *et al*., 2016) to build a time-scaled tree. Finally, we rooted the obtained timetree to the GISAID coronavirus reference genome (hCoV-19/Wuhan/WIV04/2019|EPI_ISL_402124|2019-12-30).

### Compositional segmentation algorithm

To divide the coronavirus genome sequence into an array of compositionally homogeneous, non-overlapping domains, we used a heuristic, iterative segmentation algorithm (Bernaola-Galván, Román-Roldán and Oliver, 1996; Oliver *et al*., 1999, 2004; Bernaola-Galván, Carpena and Oliver, 2008). We chose the Jensen-Shannon divergence as the divergence measure between adjacent segments, as it can be directly applied to symbolic nucleotide sequences. At each iteration, we used a significance threshold (*s* = 0.95) to split the sequence into two segments whose nucleotide composition is homogeneous at the chosen significance level, *s*. In this way, we associated a P value (P ≤ 0.05) to the final segmentation (and to the corresponding SCC value obtained, see below) of the sequence. The process continued iteratively over the new resulting segments while sufficient significance continued to appear.

### Computing the Sequence Compositional Complexity (SCC)

Once each coronavirus genome sequence was segmented at a given significance level (e.g., ≤ 0.05) into an array of statistically significant, homogeneous compositional domains, its nucleotide compositional heterogeneity was measured by computing the Sequence Compositional Complexity, or *SCC* (Román-Roldán, Bernaola-Galván and Oliver, 1998). Note that if the sequence is homogeneous, no segments should appear (SCC = 0). But if the sequence is statistically heterogeneous (at the chosen significance level), it will be segmented (SCC > 0). This is because *SCC* increased with both the number of domains in the genome and the degree of compositional differences among them. Note also that we associated a P value to each SCC value; therefore, SCC can be safely applied to any sequence size, provided the given significance threshold was reached. In this way, *SCC* is analogous to other biological complexity measures, particularly to that described by McShea and Brandon (McShea and Brandon, 2010), in which an organism is more complex if it has a greater number of parts and a higher differentiation among these parts. It should be emphasized that *SCC* is overly sensitive to any change in the RNA genome sequence, either nucleotide substitutions, indels, genome rearrangements, or recombination events, all of which could alter the number of domains or the differences in nucleotide frequencies among them. This means that a unique mutation is able to change the SCC value, but obviously as more mutations are accumulating, higher SCC values are obtained.

### Phylogenetic ridge regression

To search for trends in *SCC* values and evolutionary rates over time, phylogenetic ridge regression was applied using the *RRphylo* R package V. 2.5.8 (Castiglione *et al*., 2018). The estimated *SCC* value for each tip or node in the phylogenetic tree was regressed against its age (the phylogenetic time distance, which represents the time distance between the first sequence ever of the virus and the collection date of individual virus isolates); the regression slope was then compared to BM expectation (which models evolution according to no trend in *SCC* values and rates over time) by generating 1,000 slopes simulating BM evolution on the phylogenetic tree, using the function *search.trend* (Castiglione *et al*., 2019) in the *RRphylo* R package.

### Comparing the effects of variants on the evolutionary trend

In order to explicitly test the effect of variants and to compare variants among each other we selected 4 different trees and SCC data (s730, a727, s1871, s1990) from Table 1. In each sample, we accounted for phylogenetic uncertainty by producing 100 dichotomous versions of the initial tree by removing polytomies applying the *RRphylo* function *fix.poly* (Castiglione *et al*., 2018). This function randomly resolves polytomous clades by adding non-zero length branches to each new node and equally partitioning the evolutionary time attached to the new nodes below the dichotomized clade. Each randomly fixed tree was used to evaluate temporal trends in *SCC* and its evolutionary rates occurring on the entire tree and individual variants if present, by applying *search. trend*. Additionally, for the larger phylogenies (i.e., s1871 and s1990 lineage-wise trees) half of the tree was randomly sampled and half of the tips were removed. This way we avoided biasing the results due to different tree sizes.

## Supplementary Information

Additional details regarding the methods used in this study are provided in the Supplementary Information and in the supplemental data files available in the open repository Zenodo (https://doi.org/10.5281/zenodo.7555406).

## Acknowledgements

We gratefully acknowledge all data contributors, i.e., the Authors and their Originating laboratories responsible for obtaining the specimens, and their Submitting laboratories for generating the genetic sequence and metadata and sharing via the GISAID Initiative, on which this research is based. A complete list acknowledging all originating and submitting laboratories is available in the GISAID’s EpiCoV database (Elbe and Buckland-Merrett, 2017; Shu and McCauley, 2017; Khare *et al*., 2021) (EPI_SET_230117nx, https://doi.org/10.55876/gis8.230117nx). In the same way, we gratefully acknowledge the authors, originating and submitting laboratories of the genome sequences used for the analysis of the SARS-CoV-2 *Nextstrain* global dataset (Hadfield *et al*., 2018), downloaded on 2021-10-08; a complete acknowledgement list is shown in Supplementary Table S20 available in the open repository Zenodo (https://doi.org/10.5281/zenodo.7555406).

## Author contributions

J.L.O., M.V. and A.M. designed research; J.L.O., P.B., F.P., C.G.M, S.C., P.R., M.V. and A.M. performed research. J.L.O., P.B., F.P., C.G.M, S.C., P.R., M.V. and A.M. analyzed data; J.L.O., M.V., A.M. and P.R. drafted the paper. All authors read and approved the final manuscript.

## Funding

This project was funded by grants from the Spanish Minister of Science, Innovation and Universities (former Spanish Minister of Economy and Competitiveness) to J.L.O. (Project AGL2017-88702-C2-2-R) and A.M. (Project PID2019-105969GB-I00), a grant from Generalitat Valenciana to A.M. (Project Prometeo/2018/A/133) and co-financed by the European Regional Development Fund (ERDF). The most time-demanding computations were done on the servers of the Laboratory of Bioinformatics, Dept. of Genetics & Institute of Biotechnology, Center of Biomedical Research, 18100, Granada, Spain.

## Availability of data and materials

The data underlying this article are available in the open repository Zenodo (https://doi.org/10.5281/zenodo.7555406).

## Competing interests

The authors declare no competing interests.

## Notes

https://doi.org/10.5281/zenodo.7555406

### Competing Interest Statement

The authors have declared no competing interest.

### Summary of Updates

1. We included a most recent sample (s3037) from October 2022, thus widening the time span analyzed. 2. Figure 2 and Tables 2, 3 were updated by using this recent sample. 3. We emphasize that the samples of high-quality coronavirus genomes analyzed show sufficient variation in SCC to derive its genealogical or evolutionary relationships. 4. In the same way, we also emphasize that SCC has an associated P value, and that therefore can be safely applied to any sequence size, provided the given significance threshold was reached. 5. We now describe how the increase in the fitness of variants (by the accumulation of main and hitchhiking mutations) would lead to a change in the SCC. 6. We moved the description of timetree inference to the main text. 7. We now cite the interesting work of Cooper on the role of convergent evolution in the coronavirus.

