## Supplementary Information for "Adaptive trends of sequence compositional complexity over pandemic time in the SARS-CoV-2 coronavirus"

### Data retrieving

Random samples of coronavirus genome sequences (EPI_SET_230117nx, <https://doi.org/10.55876/gis8.230117nx>) were retrieved from the GISAID/Audacity database (Elbe and Buckland-Merrett, 2017; Shu and McCauley, 2017; Khare *et al.*, 2021). At the time of writing, this database has more than 11 million SARS‑CoV‑2 entries with complete collection date, which we used here as a proxy for the appearance over time of each strain. To retrieve accession IDs, we used both the GISAID search frontend (<https://www.epicov.org/epi3/frontend#1ba380>) and the metadata of GISAID/Audacity (<https://www.epicov.org/epi3/frontend#3bf021>). To randomly sample this huge database, we randomized the retrieved list of accession IDs, then ordering and extracting each time an elite formed by the first 500, …3,000 genome entries (Supplementary Tables S1-S19 available in Zenodo <https://doi.org/10.5281/zenodo.7555406>).

We used the quality filters provided by the GISAID website to retrieve high-quality genome sequences: only entries with complete collection date, larger than 29000 nt, with < 1% Ns and <0.05% unique amino-acid mutations (i.e., not seen in other sequences in the database) and not insertions/deletions unless verified by submitter. The virus name, the collection date (spanning from December 2019 to March 2022), and the *SCC* value for each coronavirus genome are detailed in Supplementary Tables S1‑S19 available in Zenodo (<https://doi.org/10.5281/zenodo.7555406>). An updated genomic map of the isolate Wuhan-Hu-1 (MN908947.3) we used for filtering and masking alignments of SARS-CoV-2 sequences can be found at <https://www.ncbi.nlm.nih.gov/nuccore/MN908947.3?report=graph>. Genomic information on the official reference sequence employed by GISAID (EPI_ISL_402124, hCoV-19/Wuhan/WIV04/2019, WIV04) can be found at <https://www.gisaid.org/resources/hcov-19-reference-sequence/>. We used this genome as root when inferring a phylogeny for each sample. Note that although WIV04 is twelve nucleotides shorter than Wuhan-Hu-1 at the 3' end, the two sequences are identical in practical terms, particularly the 5' UTR is the same length, and the coding regions are identical. Therefore, the coordinates and relative changes are the same whichever sequence is used (Singer *et al.*, 2020).

### Filtering and masking

We followed the steps that have been recognized so far as useful for filtering and masking alignments of SARS-CoV-2 sequences, in this way avoiding sequence oddities (Hodcroft *et al.*, 2021). First, we aligned each sample dataset to the genome sequence of the isolate Wuhan‑Hu‑1 (MN908947.3) using *MAFFT* (Katoh and Standley, 2013), following the detailed protocol at <https://virological.org/t/issues-with-sars-cov-2-sequencing-data/473>. We then mask the resulting alignment using the *Python* script ‘*mask_alignment_using_vcf.py*’ following the detailed protocols at <https://virological.org/t/masking-strategies-for-sars-cov-2-alignments/480> and <https://github.com/W-L/ProblematicSites_SARS-CoV2>. In this way, we avoid oddities in the SARS‑CoV‑2 genome sequences, such as alignment ends, which are affected by low coverage and high rate of apparent sequencing/mapping errors, recurrent or systematic sequencing errors, or homoplasic, recombination, and hypervariable sites.

### Multiple alignment and phylogeny

The masking of some sites in the first alignment with the reference sequence provokes that some of the sequences in the initial random sample become identical to others. Upon eliminating such duplicates, we realigned each sample with *MAFFT* (Katoh and Standley, 2013) but now using default options. To solve polytomies, we used the function *fix.poly* from the *RRphylo* package V.2.5.8 (Castiglione *et al.*, 2018). We then infer the best ML tree for each sample by means of the software *IQ-TREE 2* (Minh *et al.*, 2020), using the GTR nucleotide substitution model (Tavaré, 1986; Rodríguez *et al.*, 1990) and additional options suggested by the software (i.e., GTR+F+R2). We used the least square dating (LSD2) method (To *et al.*, 2016) to build a time‑scaled tree. Finally, we rooted the obtained timetree to the GISAID coronavirus reference genome (hCoV‑19/Wuhan/WIV04/2019|EPI_ISL_402124|2019-12-30).

### Sequence Compositional Complexity (SCC)

Once filtered and masked, we measured the nucleotide compositional heterogeneity of each coronavirus sequence by computing the Sequence Compositional Complexity, or *SCC* (Román-Roldán, Bernaola-Galván and Oliver, 1998). This was a two‑step process: the nucleotide sequence was first segmented into homogeneous, statistically significant compositional domains, then computing *SCC*.

#### Compositional sequence segmentation

We divided each four-symbol RNA coronavirus genome sequence into an array of compositionally homogeneous, nonoverlapping domains using a heuristic, iterative segmentation algorithm (Bernaola-Galván, Román-Roldán and Oliver, 1996; Oliver *et al.*, 1999, 2004; Bernaola-Galván, Carpena and Oliver, 2008). In brief, a sliding cursor is moved along the sequence, and the position that optimizes a proper measure of compositional divergence between the left and right parts is selected. We choose the Jensen-Shannon divergence (equations (1) and (2) in (Bernaola-Galván, Román-Roldán and Oliver, 1996)) as the divergence measure, as it can be directly applied to symbolic nucleotide sequences. If the divergence is statistically significant (at a given significance level, *s =* 0.95), the sequence is split into two segments. Note that each pair of resulting segments are more homogeneous than the original sequence. The two resulting segments are then independently subjected to a new round of segmentation. The process continues iteratively over the new segments while sufficient significance continues appearing. Since Shannon entropy is invariant under symbol interchange, the segmentation algorithm, and the *SCC* values derived from it, are invariable to sequence orientation.

Note that the statistical significance level *s* (here 0.95) is the probability that the difference between adjacent domains is not due to statistical fluctuations. By changing this parameter, one could obtain the underlying distribution of segment lengths and nucleotide compositions at distinct levels of detail (Bernaola-Galván *et al.*, 2012), thus conveniently fulfilling one of the key requirements to compute a complexity measure (Gell-Mann and Lloyd, 1996). Recent improvements to this segmentation algorithm (Bernaola-Galván *et al.*, 2012), allow segmenting sequences with long‑range correlations, as those recently reported in the coronavirus (Meraz *et al.*, 2022). Implementation details, source codes, and executable binaries for different operating systems can be downloaded from: <https://github.com/bioinfoUGR/segment> and <https://github.com/bioinfoUGR/isofinder>. The result is the segmentation of the original sequence into an array of contiguous, nonoverlapping segments (or compositional domains) whose nucleotide composition is homogeneous at the chosen significance level, *s*. As an example, the segmentation of the coronavirus reference genome is shown in Figure 1.

#### Computing SCC

Once a sequence is segmented into an array of homogeneous compositional domains at a given significance level (e.g., P value ≤ 0.05), a reliable measure of Sequence Compositional Complexity or *SCC* (Román-Roldán, Bernaola-Galván and Oliver, 1998)*,* expressed in bits by sequence position, was computed:

$$SCC=H\left( S \right)-\sum_{i=1}^{n} \frac{G_{i}}{G}H\left( S_{i} \right) [1]$$

where *S* denotes the whole genome sequence and *G* its length, *G_i_* the length of the *i*^th^ domain *S_i_*. $H\left( \cdot\right)= -\sum f{log}_{2}f$ is the Shannon entropy of the distribution of relative frequencies of symbol occurrences, *f*, in the corresponding (sub)sequence. It should be noted that the above expression is the same as the one used in the segmentation process, applying it to the tentative two new subsequences (n = 2) to be obtained in each step. Thus, the two steps of the *SCC* computation are based on the same theoretical background. Note that 1) this measure is zero if no segments are found in the sequence (the sequence is compositionally homogeneous, e.g., a random sequence) and 2) it increases/decreases with both the number of segments and the degree of compositional differences among them. In this way, the *SCC* measure is analogous to the measure used by McShea and Brandon (McShea and Brandon, 2010) to obtain complexity estimates on morphological characters: an organism is more complex if it has a greater number of parts and a higher differentiation among these parts. It should also be emphasized the high sensitivity of our measure to sequence changes. A single nucleotide substitution or one little indel suffices to alter the number, the length, or the nucleotide frequencies of the compositional domains, and therefore the resulting value for *SCC*. On the other hand, the samples of high-quality coronavirus genomes analyzed here show sufficient variation in SCC to derive its genealogical or evolutionary relationships.

### Phylogenetic ridge regression

The phylogenetic ridge regression of *SCC* was determined by using the *RRphylo* R package V. 2.5.8 (Castiglione *et al.*, 2018). In *RRphylo* the change in *SCC* value between any two consecutive tree branches aligned along a phyletic line is described by the equation *ΔSCC = β_1_1_1 +_ β_2_1_2 +_ ... _+_ β_n_1_n_* where the *β_ith_* and *l_ith_* elements represent the regression coefficient and branch length, respectively, for each *i_th_* branch along the phyletic line. The matrix solution to find the vector of *β* coefficients for all the branches is given by the equation $\hat{\beta}$ = (**L***^T^* **L** + λ**I**)^-1^ **L***^T^* *SCC*; where **L** is the matrix of tip to root distances of the tree (the branch lengths), having tips as rows, where entries are zeroes for the branches outside the tip phyletic line, and actual branch lengths for those branches along the path. *λ* is a penalization factor that avoids perfect predictions of *SCC* preventing model overfitting. The vector of ancestral states $\hat{a}$ (*SCC* values at the tree nodes) is obtained by the equation $\hat{a}=\mathbf{L}^{'}\hat{\beta}$, where $\mathbf{L}^{'}$ is the node to root path matrix, calculated as **L**, but with nodes as rows. The estimated *SCC* value for each tip or node in the phylogenetic tree is regressed against its age (the phylogenetic time distance, which represents the time distance between the first sequence ever of the virus and the collection date of individual virus isolates) and the regression slope compared to Brownian Motion (BM) expectations (which predicts no trend in *SCC* values and rates over time) by generating 1,000 slopes simulating BM evolution on the phylogenetic tree, using the function *search.trend* (Castiglione *et al.*, 2019) in the *RRphylo* R package.

### Accounting for variants diversity

In the main manuscript, we focused on early, non-variant SARS-CoV2 genomes and the three most widespread VoCs. Accounting for the potential effect of other variants requires assembling an especially large phylogenetic tree representing greater strain diversity. To this aim, we prepared a large sample, s6936 (Supplementary Table S19) with 6,936 genome sequences inclusive of six different variants (Delta: 4276, Alpha: 1323, non-variant: 1058 being overrepresented compared to others; Omicron: 44, Beta: 49). The corresponding large tree affords the possibility to test multiple variants at once and compare each of them to the general pattern of SARS-CoV2 evolution. Variants frequency changes across two orders of magnitude in the tree, from 44 (Omicron) to > 4000 (Delta). This great difference in sample size would make the regression slopes hardly comparable (given the effect size differences) and generate unmanageable computational times at addressing phylogenetic uncertainty. To solve these issues, we applied rarefaction (Sanders, 1968) to sampling size according to the equation:

$$y\left( s \right)=0.97-\left( 0.97-f_{0} \right)*e^{-f_{0}*15}$$

where $f_{0}$ is the frequency of the rarest variant (Omicron) that is 44/6936 = 6.344 10^-3^. We further reduced the frequency of non-variant genomes eliminating twenty more strains from the selection. This means the number of strains per variant changes reduces to 977. In a second experiment we rarefied according to the equation:

$$y\left( s \right)=0.91-\left( 0.91-f_{0} \right)*e^{-f_{0}*15}$$

The two sampling schemes originate the distribution of strains per variant shown in the Table S1.

Table S1. Distribution of genomes across different SARS-CoV2 strains as they are represented in the 6936 tree. The tree starts with Wuhan original samples and ends in May 2021. BV: the number of strains before the emergence of variants.

| Number of strains kept after the rarification process | | | |
| --- | --- | --- | --- |
|  | Number of kept strains | | |
|  | Full tree | Reduced tree | Extremely reduced tree |
| Omicron | 44 | 40 | 40 |
| Beta | 49 | 44 | 44 |
| Epsilon | 54 | 48 | 47 |
| Gamma | 132 | 101 | 100 |
| BV* | 1058 | 172 | 115 |
| Alpha | 1323 | 187 | 113 |
| Delta | 4276 | 385 | 129 |
| *BV represent the number of strains before the emergence of variants. | | | |

#### SCC trends over time

There is significant and negative temporal trend in SCC for the 977 lineages trees in 65 out of 100 replications. All individual variants show substantial support for significant and negative trends in *SCC* values (Table S2). Comparing the slope of *SCC* versus time regression of individual variants, we found substantial support for the slope of variants being significantly steeper than the slope (intensity of the trend) of the rest of the tree (Table S2).

Table S2. Percentages of significant results of *SCC* and *SCC* evolutionary rates versus time regressions performed on 100 randomly sampled 977 lineage-wide phylogenetic trees. % slope = percentage of significant regression slopes; % slope difference = percentage of significant regression slope differences.

|  |  | **SCC values** | **SCC evolutionary rates** |
| --- | --- | --- | --- |
|  |  | **% Slope** | **% Slope difference** |
| Alpha | positive | 0 | 0 |
|  | negative | 100 | 3 |
| Beta | positive | 0 | 0 |
|  | negative | 100 | 0 |
| Delta | positive | 0 | 13 |
|  | negative | 94 | 27 |
| Epsilon | positive | 0 | 0 |
|  | negative | 100 | 0 |
| Gamma | positive | 2 | 0 |
|  | negative | 83 | 0 |
| Omicron | positive | 0 | 0 |
|  | negative | 97 | 74 |

We found a significant and positive trend in *SCC* evolutionary rates through time in 60% cases. The slope of *SCC* rates through time regression for Omicron is significantly lower (more negative) than the slope computed for rest of the tree (Table S2) in 74% of the cases. The same figure for Delta is significantly higher in 13% and significantly lower in 27% cases compared to the resto of the tree. Other variants show low to no support for either trend (Table S2). The phylogenetic ridge regressions for SCC and its evolutionary rate on the s6936 sample reduced to 977 lineages are shown in Figure S1.

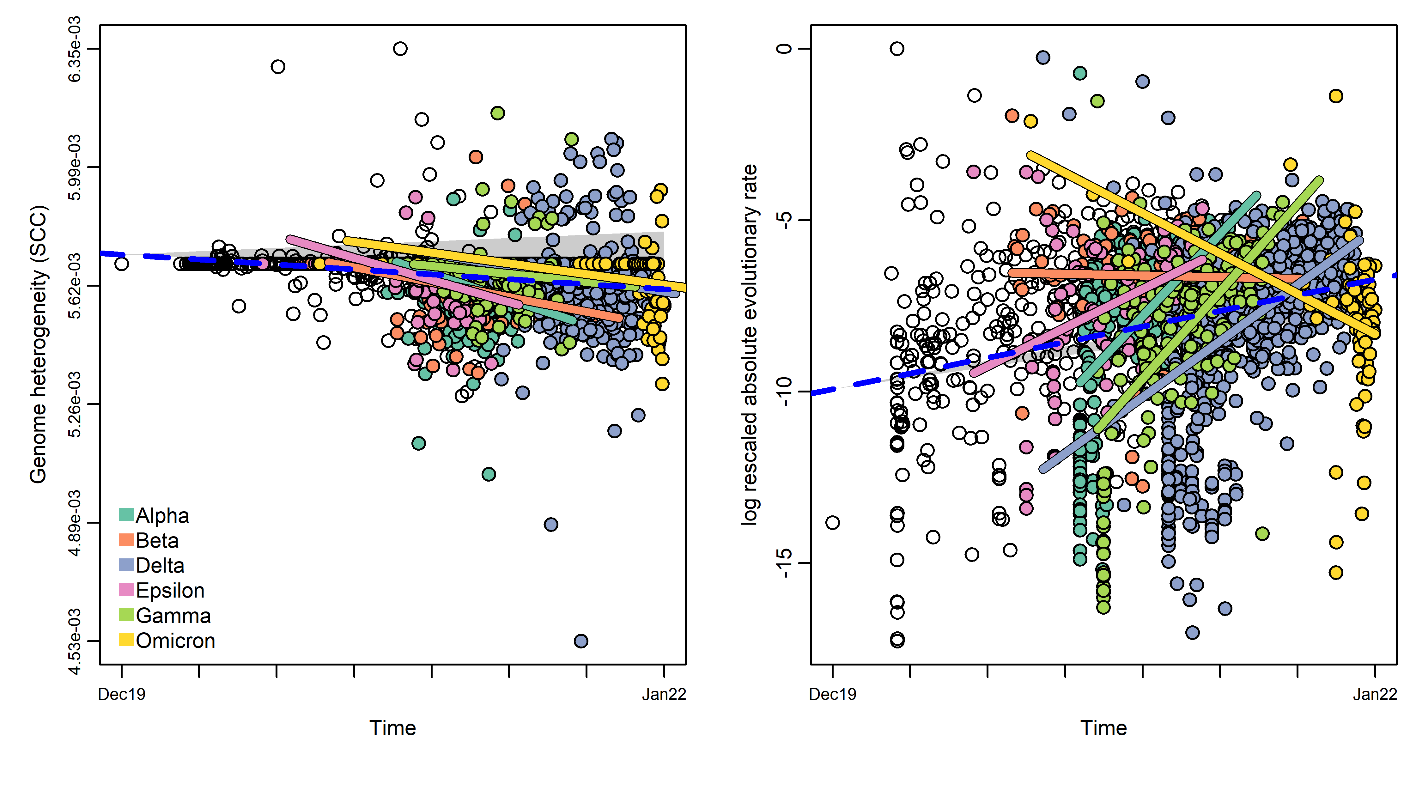

Figure S1. Phylogenetic ridge regressions for SCC (left) and its evolutionary rate (right) as detected by the *RRphylo* R package (Castiglione *et al.*, 2018) on the s6936 sample reduced to 977 lineages. For SCC, the estimated value for each tip in the phylogenetic tree is regressed (dashed blue line) against its age (the phylogenetic time distance, meant as the collection date of each virus isolate). The rescaled evolutionary rate was obtained by rescaling the absolute rate in the 0-1 range and then transforming to logs to compare to the BM expectation. The statistical significance of the ridge regression slopes was tested against 1,000 slopes obtained after simulating a simple Brownian evolution of the SCC in the phylogenetic tree. The 95% confidence intervals around each point produced according to the BM model of evolution are shown as shaded areas. Dots are colored according to the variant they belong or left blank for strains collected before the appearance of variants.

Repeating the same analysis with 588-lineages trees and with the trees obtained drawing variant sequences randomly give the same exact insights, showing a neat temporal trend for decrease in SARS-CoV2 SCC and accelerated rates of evolution, especially clear in the Omicron variant. There is significant and negative temporal trend in SCC for the full tree (notice, ‘full’ here means 588 lineages randomly extracted from the 6936 tree) in 73 out of 100 randomly sampled trees. All individual variants show substantial support for significant and negative trends in *SCC* values (Table S3).

Table S3. Percentages of significant results of SCC and SCC evolutionary rates versus time regressions performed on 100 randomly sampled 589 lineage-wide phylogenetic trees. % slope = percentage of significant regression slopes; % emmeans difference = percentage of significant estimated marginal mean difference; % slope difference = percentage of significant regression slope differences.

|  |  | **SCC values** | **SCC evolutionary rates** |
| --- | --- | --- | --- |
|  |  | **% Slope** | **% Slope difference** |
| Alpha | positive | 0 | 0 |
|  | negative | 99 | 3 |
| Beta | positive | 0 | 0 |
|  | negative | 99 | 0 |
| Delta | positive | 1 | 0 |
|  | negative | 86 | 9 |
| Epsilon | positive | 0 | 0 |
|  | negative | 100 | 0 |
| Gamma | positive | 6 | 0 |
|  | negative | 77 | 0 |
| Omicron | positive | 0 | 0 |
|  | negative | 98 | 88 |

We found a significant and positive trend in SCC evolutionary rates through time in 55% cases. The slope of SCC rates through time regression for Omicron is significantly lower than the slope computed for rest of the tree (Table S3) in 88%. Other variants show low to no support for a significant trend in rates (Table S3). The phylogenetic ridge regressions for SCC and its evolutionary rate on the s6936 sample reduced to 588 lineages are shown in Figure S2.

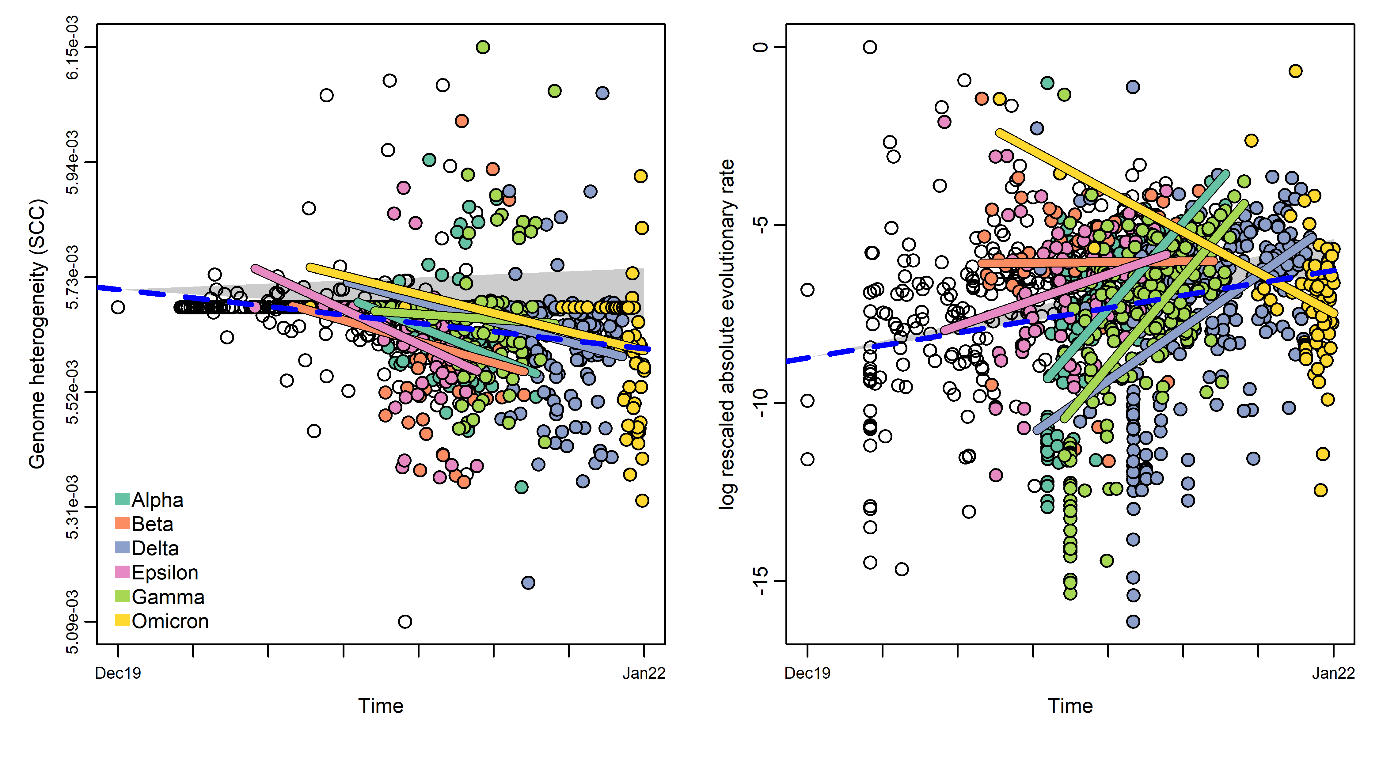

Figure S2. Phylogenetic ridge regressions for SCC (left) and its evolutionary rate (right) as detected by the *RRphylo* R package (Castiglione *et al.*, 2018) on the s6936 sample reduced to 588 lineages. For SCC, the estimated value for each tip in the phylogenetic tree is regressed (dashed blue line) against its age (the phylogenetic time distance, meant as the collection date of each virus isolate). The rescaled evolutionary rate was obtained by rescaling the absolute rate in the 0-1 range and then transforming to logs to compare to the BM expectation. The statistical significance of the ridge regression slopes was tested against 1,000 slopes obtained after simulating a simple Brownian evolution of the SCC in the phylogenetic tree. The 95% confidence intervals around each point produced according to the BM model of evolution are shown as shaded areas. Dots are colored according to the variant they belong or left blank for strains collected before the appearance of variants.

### Checking results reliability

Despite the quality filters of the GISAID website (Elbe and Buckland-Merrett, 2017; Shu and McCauley, 2017; Khare *et al.*, 2021) used to retrieve high‑quality genome sequences, the steps we followed for filtering and masking the sequences, as well as the relative large size of the random samples chosen for the analysis (Table 1 and Supplementary Tables S1-S19, some concerns could be raised on the reliability of our results, mainly on the algorithms we used for multiple alignment or to infer phylogenetic trees.

To check for the reliability of our results when the multiple alignment tool is changed, we repeated the analysis of the s987 sample using *Nextalign* (Aksamentov *et al.*, 2021) instead of *MAFFT* (Katoh and Standley, 2013). *Nextalign* was a codon-aware pairwise sequence aligner for similar viral genomes, which allows unambiguous calling of amino-acid changes associated with changes in the nucleotide sequence. Using this aligner, we also found in this sample a highly‑significant decreasing trend for SCC (slope = -59.8, P value ≤ 0.002), as well as an increasing trend for the evolutionary rate (slope = 16316, P value ≤ 0.001).

In the same way, we repeated our analyses using coronavirus samples and trees obtained by other research groups, finding qualitatively comparable results. An example was the analysis we made of the SARS-CoV-2 *Nextstrain* global dataset (Hadfield *et al.*, 2018) containing 3,059 genomes downloaded from <https://nextstrain.org/ncov/open/global?f_host=Homo%20sapiens> on 2021-10-08. The sequences for this dataset (in Fasta format) and the ML phylodynamic tree obtained by these authors by means of the TreeTime software (Sagulenko, Puller and Neher, 2018) are available at Zenodo (<https://doi.org/10.5281/zenodo.7555406>). We obtained a decreasing, although marginally significant trend, for SCC (slope = ‑0.01, P value ≤ 0.122), as well as an increasing, highly significant trend for the evolutionary rate (slope = 0.89, P value ≤ 0.001).

### Supplemental files

The supplementary data files underlying this article are available in the open repository Zenodo (<https://doi.org/10.5281/zenodo.7555406>):

| **File** | **Description** |
| --- | --- |
| SupplementaryTables S1-S19.zip | Excel supplementary tables: The strain name, the collection date, and the SCC values for each analyzed genome. |
| nextstrain_ncov_open_global_timetree.nwk | ML phylodynamic tree for the Nextstrain sample |
| SupplementaryTable S20.pdf | A complete list acknowledging the authors, originating and submitting laboratories of the genetic sequences we used for the analysis of the Nextstrain sample. |
| Nextstrain_sample_fasta_3059.zip | Nextstrain sample (sequences in Fasta format) |
| PhylogeneticTimetrees_NewickFormat.zip | Phylogenetic timetrees (Newick format). |
